# Beyond Prior and Volatility: The Distinct Iterative Updating Account of ASD

**DOI:** 10.1101/2022.01.21.477218

**Authors:** Zhuanghua Shi, Fredrik Allenmark, Laura A. Theisinger, Rasmus L. Pistorius, Stefan Glasauer, Hermann J. Müller, Christine M. Falter-Wagner

**Author notes:** Corresponding authors: Dr. Zhuanghua Shi, Dr. Christine M. Falter-Wagner. **Author’s contributions.** Z.S., F.A., and C.F. study conceptualization; L.T., C.F., S.G. funding acquisition; C.F., L.T., R.P., and F.A. participant recruitment; Z.S., F.A., and C.F. supervision; L.T., R.P., F.A. data collection; L.T., R.P., F.A., S.G., and Z.S. data analysis and manuscript draft; Z.S. and S.G. computational modeling; Z.S., L.F., F.A., R.P., S.G., H.M., and C.F. critical revision and editing. **Competing interests.** The authors declare no competing financial interests. **Classification**: Social Science - Psychological and Cognitive Sciences.

## Abstract

The nature of predictive-processing differences between individuals with autism spectrum disorder (ASD) and typically developing (TD) individuals is widely debated. Some studies suggest impairments in predictive processing in ASD, while others report intact processes, albeit with atypical learning dynamics. Here, we assessed duration reproduction tasks in high- and low-volatility settings to examine the updating dynamics of prior beliefs and sensory estimates. Employing a two-state Bayesian model, we differentiated how individuals with ASD and TD controls update their priors and perceptual estimates, and how these updates affect long-term prediction and behavior. Our findings indicate that individuals with ASD use prior knowledge and sensory input similarly to TD controls in perceptual estimates. However, they place a greater weight on sensory inputs specifically for iteratively updating their priors. This distinct approach to prior updating led to slower adaptation across trials; individuals with ASD relied less on their priors in perceptual estimates during the first half of sessions but achieved comparable integration weights as TD controls by the end of the session. By differentiating these aspects, our study highlights the importance of considering inter-trial updating dynamics to reconcile diverse findings of predictive processing in ASD. In consequence to the current findings, we suggest the distinct iterative updating account of predictive processing in ASD.

**Significance Statement:** Research on predictive processing in Autism Spectrum Disorder (ASD) remains controversial. The current study employed a two-state Bayesian model in varied volatility settings to explore inter-trial updating dynamics in ASD compared to typically developing (TD) peers. We found that individuals with ASD, while utilizing prior knowledge similarly to TD controls, place a disproportionate emphasis on sensory inputs when updating their priors. This unique pattern of slower adaptation during iterative updating leads to significant behavioral differences in the first half of trials between the two groups, but comparable levels by the end of the session. These findings not only highlight the importance of considering different timescales and dynamic updating processes in ASD, but also suggest that the predictive processing framework in ASD involves unique prior updating mechanisms that is likely associated with increased sensory reliance.

## Introduction

Autism Spectrum Disorder (ASD) manifests as challenges in social interaction, communication, and repetitive behaviors. Individuals with ASD often exhibit heightened sensory sensitivity and adaptation difficulties (Gernert et al., 2020; Leekam et al., 2007; Tomchek & Dunn, 2007).

Predictive processing and Bayesian inference provide promising frameworks for understanding the autistic cognitive profile, pointing to a particular style of processing sensory information and prior knowledge in ASD (Cannon et al., 2021; Lawson et al., 2014, 2017; Noel et al., 2022; Noel & Angelaki, 2023; Palmer et al., 2017; Pellicano & Burr, 2012; Sinha et al., 2014; Van de Cruys et al., 2014). Bayesian inference posits that perception and action result from the constant fine-tuning of predictions and minimization of prediction errors by combining sensory input with previously acquired knowledge (‘priors’), weighted by their associated uncertainties. While the theories put forward thus far agree that predictive processing is different in ASD, they disagree on the specific mechanistic aspects producing the impairment, which can be broadly subsumed into two kinds of model. The “attenuated-prior” model argues that individuals with ASD have weaker priors, resulting in a reduced reliance on the priors in making predictions (Karaminis et al., 2016; Pellicano & Burr, 2012; Turi et al., 2015). Conversely, according to the “heightened and inflexible precision of prediction errors in Autism” (HIPPEA) theory, the sensory atypicalities in ASD are attributable to an overemphasis of prediction errors (Van de Cruys et al., 2014). Lawson and colleagues (2014) offer a similar perspective, highlighting an impaired ability in ASD to downweight sensory input, potentially tipping the balance towards sensory input over prior beliefs for the final perceptual decisions. Whatever the presumed cause of the atypicality, both types of model agree that predictive processing is generally different in ASD (Sinha et al., 2014).

Contrary to the view that individuals with ASD form compromised prior beliefs, many types of prior – particularly those deriving from experience or top-down knowledge – actually remain intact (Angeletos Chrysaitis & Seriès, 2023). For instance, individuals with ASD can accurately form one-shot priors, such as recognizing a Dalmatian dog camouflaged by black patches (Van de Cruys et al., 2018), and show typical performance in tasks related to the processing of gaze direction (Pantelis & Kennedy, 2017; Pell et al., 2016), statistical learning of likely distractor locations (Allenmark et al., 2021), and using external reference coordinates in tactile spatial tasks (Hense et al., 2019). These observations have led Palmer and colleagues (2017) to challenge the adequacy of simplistic Bayesian models that only account for the integration of sensory input with a single prior. Noel and Angelaki (2023) recently advocated examining ASD across different descriptive levels, including causal inference at the computational level (Noel et al., 2022), divisive normalization at the algorithmic level, and excitatory-to-inhibitory imbalance at the neural-implementation level (Rosenberg et al., 2015), yielding a more refined picture of phenotypes of ASD.

Of note, many relevant studies of ASD have focused on interpreting averaged outcomes as representing the ‘percept’ and so have disregarded a potentially important aspect: that of short-term, iterative and dynamic updating of priors from one trial to the next. In line with the latter, recent reports indicate that individuals with ASD may be less influenced by preceding events (Lieder et al., 2019) and may adapt more slowly to changes (Vishne et al., 2021) compared to their matched peers. Importantly, such distinctive patterns in dynamic updating might not necessarily compromise the formation of long-term priors, but they could impact average outcomes across different time scales. Moreover, existing accounts do not explicitly distinguish the dynamic updating of perceptual estimates from dynamic updating of the prior. Here, we propose that a direct investigation of how individuals with ASD handle short-term updates and adjust to long-term environmental changes through dynamic iterative Bayesian updating modeling can resolve some discrepancies in the literature and enhance our understanding of the mechanisms underlying atypical responses in ASD.

Accordingly, we employed a duration-reproduction task with varied sequential volatility (Glasauer & Shi, 2021) to differentiate between short-term-prior and perceptual-estimate updating and long-term integration. The reproduction task is widely recognized for its effectiveness in assessing the integration of sensory measurements with priors, thus facilitating the assessment of integration weights within a Bayesian framework (Cicchini et al., 2012; Jazayeri & Shadlen, 2010; Karaminis et al., 2016; Shi, Ganzenmüller, et al., 2013). It also supports the application of dynamic-updating models (Glasauer & Shi, 2021, 2022; Petzschner & Glasauer, 2011). Our approach, inspired by recent findings (Glasauer & Shi, 2022), assumes that the updating of priors varies based on expected trial-to-trial variability or volatility. So, across two sessions, we maintained the same set of durations while varying their sequential presentation volatility, thereby isolating the effects of short-term trial-wise perceptual-estimate updating (local volatility) from iterative updating of the long-term prior. Crucially, we employed iterative two-state Bayesian modeling (Glasauer & Shi, 2022), which tracks both the integration of sensory inputs with priors and the dynamic updating of these priors while clearly delineating them from the separate influences of the updating of perceptual estimates.

## Results

We asked participants (ASD, *n* = 32; TD, *n* = 32) to observe a visual disk that appeared for a certain time duration, ranging from 100 to 1900 ms (see *Methods*), and then to replicate the observed duration by pressing a key for the requisite time (Figure 1a). This task was completed across two separate sessions: while the same set of durations was presented in each session, the volatility - or unpredictability - of the presented durations from one trial to the next differed between the two sessions (depicted in Figure 1b). Integrating prior knowledgeof the sample distribution of these durations typically induces a strong central-tendency effect, with shorter durations tending to be overestimated and longer ones underestimated (Cicchini et al., 2012; Jazayeri & Shadlen, 2010; Lejeune & Wearden, 2009; Shi, Church, et al., 2013). By assessing this central-tendency effect, we can estimate the weight of the prior in the integration process.

**Figure 1.**
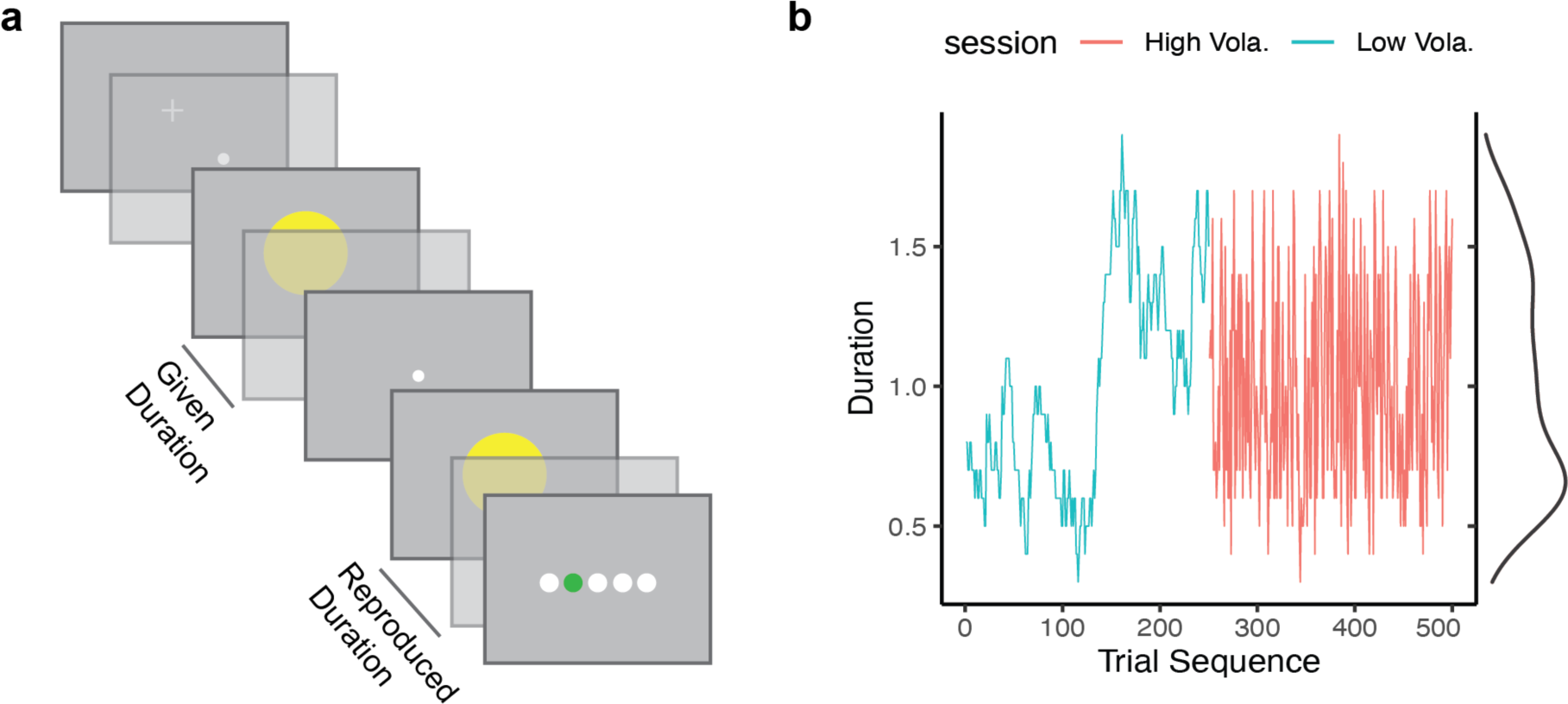
**(a)** Schematic illustration of a trial sequence used in the duration-reproduction task. (**b**) Example duration (trial) sequences in two consecutively performed ‘duration-*volatility*’ sessions. The session depicted on the left (in cyan) presents a low-volatility sequence (Low Vola.), and the session depicted on the right (in red) a high-volatility sequence (High Vola.). Both sessions presented exactly the same durations (i.e., the same density function depicted on the right), differing only in their presentation orders.

### Central-tendency Effect and Impact of Volatility

To assess the central tendency effect quantitatively, we first pre-processed each dataset of reproduced durations by removing outliers, defined as reproduced durations outside the range of [Duration/3, 3×Duration]. Such extreme reproductions, occurring on only 0.58% of all trials, were excluded from further analysis. Given the linear nature typically associated with the central-tendency effect (Glasauer & Shi, 2021; Shi, Church, et al., 2013), we used linear regression to estimate this effect. As depicted in Figures 2a and 2b, both the ASD and TD groups exhibited a linear decrease in reproduction error with increasing duration – a pattern consistent across both the low- and high-volatility sessions. Generally, shorter durations tended to be overestimated and longer durations underestimated. A steeper linear trend - indicative of a stronger central-tendency bias - was observed in the high volatility session.

**Figure 2.**
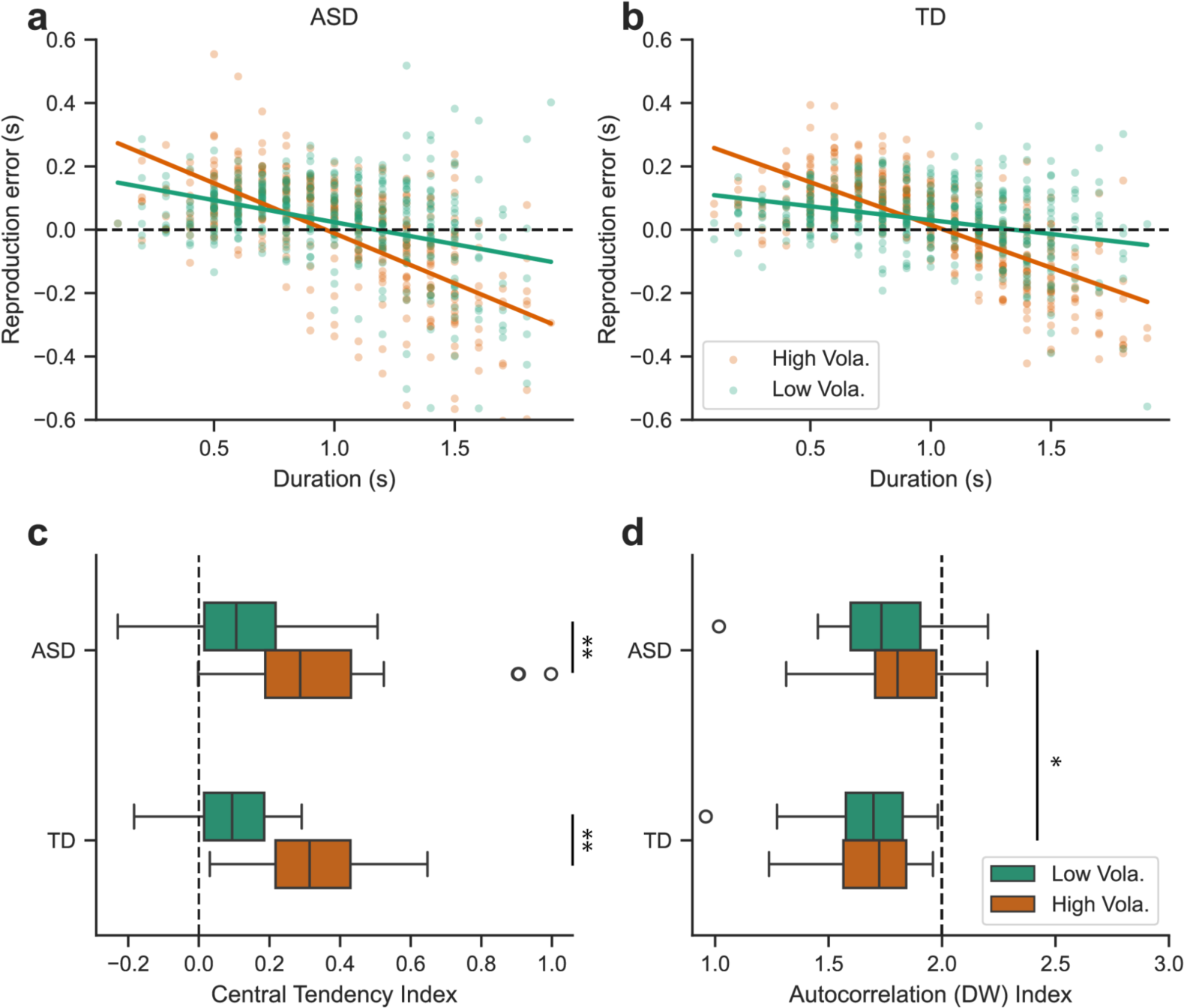
Average reproduction error as a function of the (to-be-reproduced) stimulus duration for the ASD (**a**) and TD (**b**) groups (each group *n*=32), separately for the high (red) and low (green) volatility sessions. Each individual dot stands for a single participant’s mean reproduction error for that given duration. Dots above the horizontal dashed line indicate overestimations and dots below the line underestimations, respectively. Solid lines indicate the fitted trend of the reproduction errors. Notably, the steepness of the fitted trends is higher in the high-vs. the low-volatility session, indicative of stronger central-tendency biases during high-volatility conditions. (**c**) Boxplots of the Central-Tendency Index (CTI) for both the ASD and TD groups, separately for the high- and low-volatility sessions. A CTI of 0 implies a lack of a central-tendency effect, whereas a CTI of 1 indicates a pronounced central-tendency effect. The left and right boundaries of the box denote the interquartile range (IQR), from the 25th to the 75th percentile, while the whiskers indicating 1.5 times the IQR. Three outliers were observed in the ASD group during the high-volatility session. (**d**) The boxplots of the average Durbin-Watson (DW) autocorrelation indices, separately for the high- and low-volatility sessions, for both the ASD and TD groups. The dashed line, representing a DW value of 2.0, signifies the absence of autocorrelation, while a DW value less than 2 indicates a positive autocorrelation. * denotes *p* < .05, ** *p* < .01.

We quantified the weight of the prior in sensory-prior integration using the negative slope of the regression line, termed the central-tendency index (CTI). A CTI of 0 signifies the absence of a central tendency effect, while a CTI of 1 indicates a pronounced central tendency bias. Figure 2c presents boxplots of the CTIs, differentiated by Volatility and Group. A Volatility-by-Group mixed-design ANOVA on the CTIs yielded a robust main effect of *Volatility*, *F*(1,62) = 80.11, *p* < .001, η*_p_*^2^ = .56, with no significant effects for *Group* or the *Volatility × Group* interaction (*F*s < 0.23, *p*s > .63).

In the ASD group, three ‘outlier’ participants exhibited a substantial central-tendency effect (CTI > 0.9). Essentially, these participants reproduced similar durations across all tested durations in the high-volatility session, indicative of an over-reliance on prior knowledge, giving rise to a remarkable level of rigidity in their responses (see supplementary plot in Appendix 1 for more detail). Excluding these three outliers and their matched controls did not change the pattern of statistical results (*Volatility*: *F*(1, 56) = 81.37, *p* < .001, η*_p_*^2^ =.59; *Group*, *F*(1, 56) = 1.734, *p* = .193, η*_p_*^2^=.03; *Volatility × Group* interaction, *F*(1, 56) = 3.324, *p* = .074, η*_p_*^2^ =.056).

Moreover, the average reproduction errors skewed towards overestimation. A mixed-design ANOVA of the errors with the factors Volatility (within-participant) and Group (between-participant) revealed a significant main effect of Volatility, *F*(1,62) = 5.57, *p* = .021, η^2^_p_=.08: participants overestimated durations more in the low-volatility session (by 39 ms) than in the high-volatility session (by 22 ms). Neither the Group factor nor the interaction between Group and Volatility were significant, *F*s < 0.698, *p*s > .4, *η_p_*^2^s < .01.

### Auto-correlation Reveals a Reduction of Positive Correlation in ASD

If reproduction errors were solely driven by the central tendency effect, the residuals from the regression analysis - the differences between the data and regression line - should represent independent random errors. However, our analysis of the residuals revealed a marked differential autocorrelation, showing a dependency of the reproduction error on the current trial (*n*) based on the error of the immediately preceding trial (*n*–1). Figure 2d shows the average Durbin-Watson (DW) statistics from the regression analysis for the two groups, separately for the two sessions. The DW statistic is an indicator of the presence of autocorrelation at lag 1 (correlation of trial *n* with trial *n*–1), which ranges from 0 to 4. A DW value of 2 suggests no autocorrelation, while values below 2 indicate positive autocorrelation (with a DW of zero being perfect positive autocorrelation).

A mixed-design ANOVA revealed the ASD group to display a significantly less positive autocorrelation at lag 1 compared to the TD group, *F*(1,62) = 4.50, *p* = .038, *η_p_*^2^ =.07 (excluding outliers: *F*(1,56) = 8.36, *p* = .005). The DW scores for both groups were significantly below 2 (ASD: 1.78, *t*(63) = −7.9, *p* < .001; TD: 1.68, *t*(63) = −11.9, *p* < .001). The positive autocorrelations together with the group difference indicate that the central tendency alone does not fully capture the differences between the two groups, in particular as regards the inter-trial dependencies. However, the main effect of *Volatility* and the *Volatility × Group* interaction failed to reach significance (*F*s < 2.19, *p*s > .145).

### Distinct Prior Updating Revealed by Two-State Iterative Model

The autoregression analysis indicated that the notable differences between the two groups likely stemmed from variations in short-term inter-trial updating. To more closely model these dynamics, we employed Glasauer and Shi’s two-state model (Glasauer & Shi, 2022): a hybrid Bayesian iterative-updating model designed to capture both the long-term central-tendency effects and short-term inter-trial sequential dependencies. This model posits that an individual’s internal prior belief or prediction, represented by a *prior* distribution, is malleable and evolves through iterative integration of new sensory information, represented by the *likelihood* distribution^1^. This integration results in a revised belief, the *posterior* distribution. From this posterior distribution, one can compute an estimate by applying a certain cost function, such as the maximum a-posteriori probability estimate (MAP). Unlike simplistic trial-independent Bayesian estimation, the two-state model allows the mean of the prior distribution to vary across trials, rather than being a static value. In essence, each new prediction for the subsequent trial is derived from the current posterior distribution (Figure 3) by applying knowledge about how the state of the world develops over time. This two-state model can be formulated as a Kalman filter process (Glasauer & Shi, 2022).

**Figure 3.**
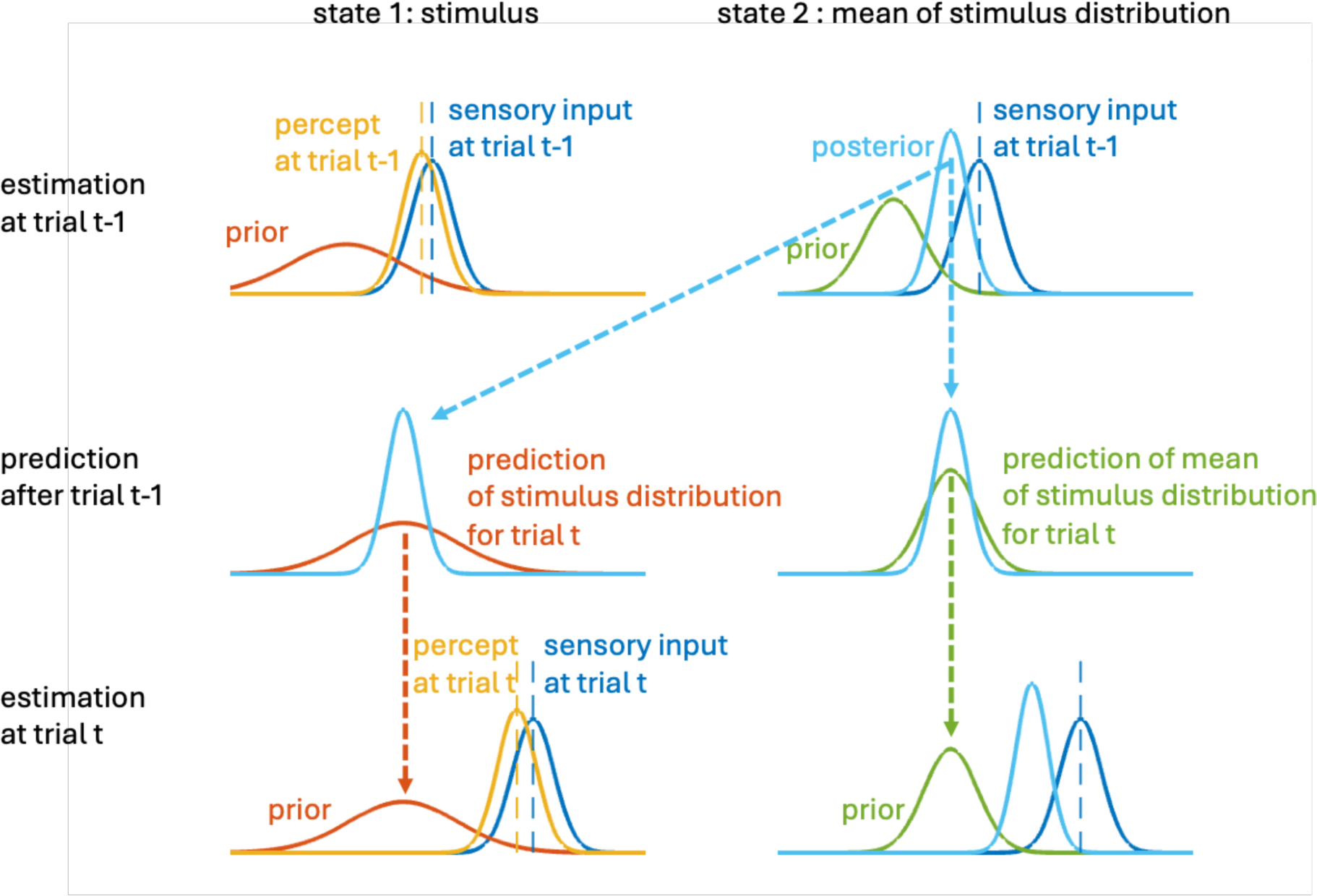
Two-state iterative Bayesian updating model. The **left** panels illustrate the iterative updates of the perceptual estimate of the stimulus (State 1), while the **right** panels depict the iterative updates of the mean of stimulus distribution (State 2). On trial *t-1* (**top row**), the percept (yellow) is computed by multiplying the prior of the stimulus (red) and the sensory likelihood (dark blue), which corresponds to Eq. (4). The posterior distribution (light blue) of the stimulus mean is derived by combining the prior of the mean distribution (green) with the sensory likelihood (dark blue), which corresponds to Eq. (5). After trial *t-1* (**middle row**), the stimulus mean distribution (light blue) for trial *t* is updated from the posterior distribution (indicated by the dashed blue arrow). The stimulus prior (red) for trial *t* is then formulated for the prediction. Additionally, the prediction of the mean of stimulus distribution (green) for trial *t* is computed from the posterior distribution (light blue). On trial *t* (**bottom row**), these processes are repeated as in trial *t-1*.

The generative process describes how the two states of the model, the stimulus duration (*x_i_*) and the mean of the stimulus distribution (*m_i_*) on the current trial *i*, evolve over time:

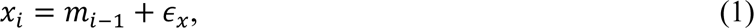

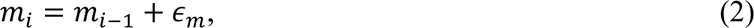

where ɛ_*x*_ and ɛ_*m*_ are normally distributed noise with variances *v* and *q*, respectively. Thus, it is assumed that the mean of stimulus distribution *m_i_* changes by a random amount each trial, and that the actual stimulus *x_i_* is drawn from that distribution. The actual sensory measurement *z_i_* is the stimulus (*x_i_*), corrupted by the sensory noise η ∼ *N(0, r)*:

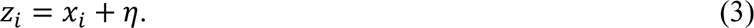

The estimate of the two states can be written as:

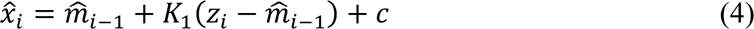

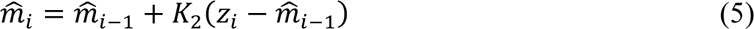

where *K_1_* and *K_2_* are the Kalman gains to determine the estimates of the current duration *x_i_* and the mean of the stimulus distribution 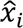, respectively, and *c* is a constant bias. The Kalman gain *K_i_* is essentially a weighting factor that determines how much weight is assigned to the new prediction error 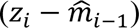. If the Kalman gain is high, the system places more trust on the new sensory information; if it is low, the system places greater trust on the prediction (prior).

To ensure that the two-state model performs well for the behavioral data, we fitted the model with three parameters: the variance ratio *v/r*, the variance ratio *q/r*, and the constant bias *c*. The model outperformed the standard linear regression, reducing the Akaike Information Criterion (AIC) by 10.75, which indicates a markedly better fit to the behavioral data. Figure 4 shows the mean estimates for two Kalman gains, *K_1_* and *K_2_*, along with the general bias (*c*). On average, *K_1_* and *K_2_* were lower in more predictable (low-volatility) environments, while the general bias (*c*) was elevated. Additionally, for the ASD group, *K_2_* was numerically higher and the bias (*c*) was numerically smaller compared to the TD group.

**Figure 4.**
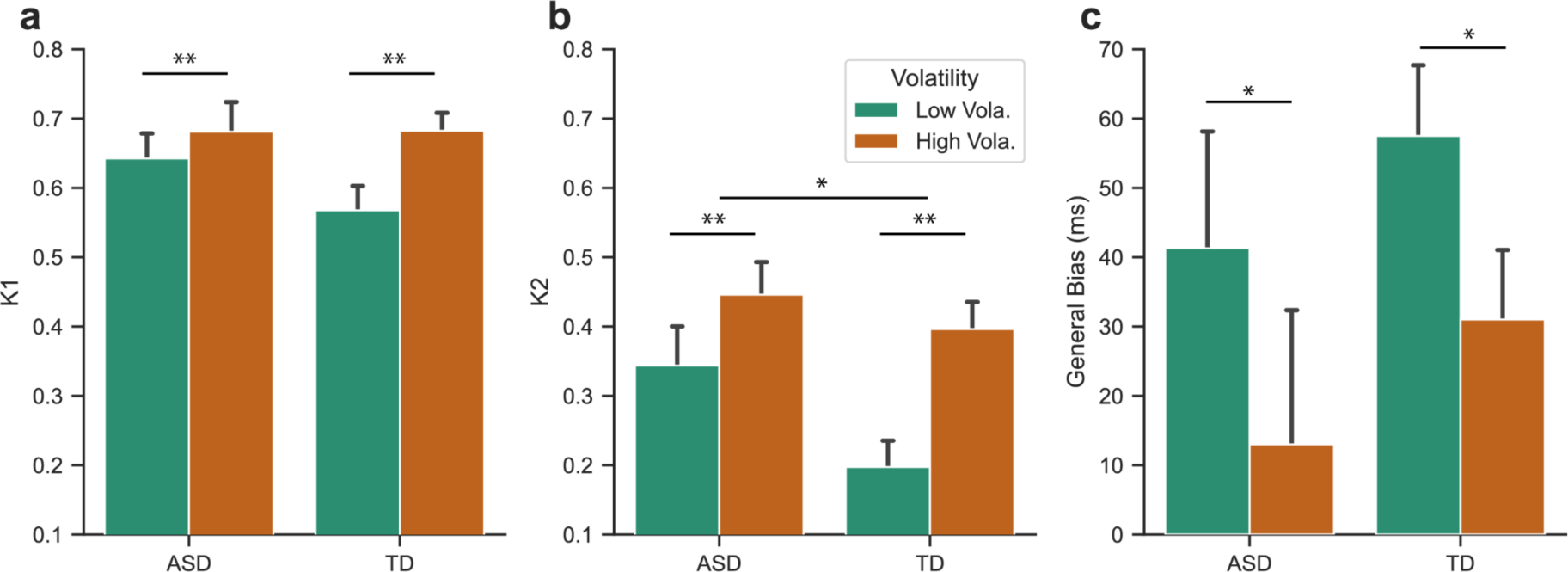
(**a**) Estimated Kalman gain *K_1_* for the state of duration prediction; (**b**) estimated Kalman gain *K_2_* for the updating of the prior belief; (**c**) general overestimation (ms), separately for two groups (ASD vs. TD) and volatility sessions (Low vs. High). Each group *n*=32. Error bars denote one standard error. * denotes *p* < .05, ** *p* < .01.

A mixed-design ANOVA of the Kalman gain *K_1_*, with *Group* and *Volatility* as factors, yielded only a significant *Volatility* effect, *F*(1, 62) = 10.61, *p* = 0.002, η*_p_*^2^=.146: the weight *K_1_* was higher in the high-(68.2%) vs. the low-volatility (60.5%) session, that is: participants relied more on the sensory input rather than their prior when making estimates in the high-volatility session. There were no main or interaction effects involving *Group*, *F*s < 2.62, *p*s > .11, η*_p_*^2^s< .04.

Conversely, a similar analysis of the Kalman gain *K_2_* revealed a significant *Group* difference, *F*(1,62) = 4.23, *p* = 0.04, η*_p_*^2^=.146: on average, individuals with ASD exhibited a higher K_2_ (0.395) than their TD counterparts (0.297), indicative of a stronger tendency of the former to revise their priors. *Volatility* also had a significant effect on K_2_, *F*(1,62) = 11.75, *p* = 0.001, η*_p_*^2^=.159: participants were more inclined to update their priors (i.e., less reliable prediction), in the high-(0.42) vs. the low-volatility (0.27) environment. The *Group × Volatility* interaction, however, was non-significant, *F*(1,62) = 1.21, *p* = 0.276, η*_p_*^2^=.019.

Another mixed-design ANOVA revealed *Volatility* to be the only significant factor affecting the general bias, *F*(1, 62) = 11.83, *p* = 0.001, η*_p_*^2^=.16, corroborating earlier reported behavioral results (there were no – main and/or interaction – effects involving *Group*, *F*s < 0.794, *p*s > .37, *η_p_*^2^s < .013).

In summary, while *Volatility* impacted all three parameters examined, a differential effect of the ASD vs. the TD group was observed only in the Kalman gain *K_2_*.

### First vs. Second Half: Impact of Updating Parameters on the Weight of the Prior

To further investigate whether greater emphasis on sensory input in short-term belief updating, as indicated by a higher Kalman gain *K_2_*, affects the speed of adjusting the long-term prior, we analyzed the first half of trials in each session. We hypothesized that if individuals with ASD start with placing a low weight on the prior, a reduced central-tendency effect should be observable early on in the session (here tested in the first half of trials). We further hypothesized that if individuals with ASD update their prior only slowly, their central-tendency effect should eventually become comparable to that of TD individual (in the second half of the trials).

Given that we only applied the first half of the trials (limiting data quality), we excluded the three ‘outlier’ individuals and their counterparts from this analysis. Figure 5a shows the central-tendency effect from these initial trials. A mixed-design ANOVA of the CTIs revealed significant main effects of both *Group* (*F*(1, 56) = 6.236, *p* = .015, *η_p_^2^*=.1) and *Volatility* (*F*(1, 56) = 24.74, *p* < .001, *η_p_^2^*=.31) – which is in contrast to the non-significant main effect of Group when tested across all trials, even when excluding the ‘outlier’ individuals. The central-tendency bias was significantly smaller for the ASD group than for the TD group in these initial trials (while being comparable across all trials), indicating slower adaptation in individuals with ASD. Similar to the analysis of all trials, the Durbin-Watson (DW) values were significantly higher for the ASD group compared to the TD group already in the first half of trials, *F*(1, 56) = 7.2, *p* = .01, *η_p_^2^*=.114 (Figure 5b).

**Figure 5.**
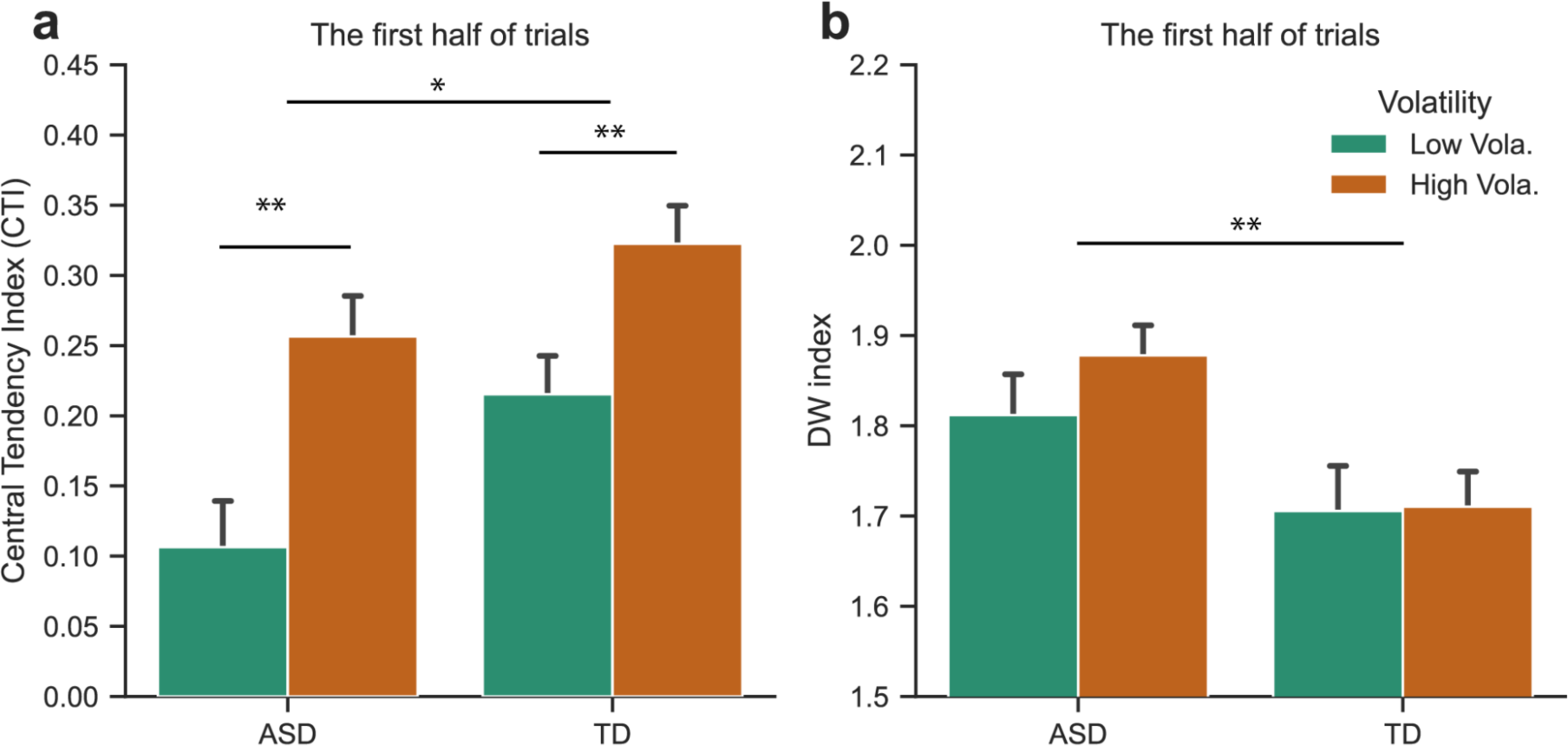
(**a**) Average central-tendency indices (CTI), and associated standard errors, for the ASD and TD groups (each group *n*=29), separately for the high- and low-volatility sessions. A CTI of 0 implies a lack of a central-tendency effect, whereas a CTI of 1 indicates a pronounced central-tendency effect. (**b**) Average Durbin-Watson (DW) autocorrelation indices, and associated standard errors, separately for the high- and low-volatility sessions, for both the ASD and TD groups. A DW value of 2.0 signifies the absence of autocorrelation, while a DW value less than 2 indicates a positive autocorrelation. Error bars denote one standard error. * denotes *p* < .05, ** *p* < .01.

Next, we re-estimated the Kalman gains *K_1_* and *K_2_* based on the first half of trials in each session and compared them to the parameters obtained for the entire set of trials. Interestingly, *K_2_* remained relatively stable across analyses of the first half of trials versus the entire set of trials, with variations within 2.2% across all conditions and groups (*F*s < 0.074, *p*s > 0.74). However, there were already significant differences in *K_2_* between the two groups (*Group*, *F*(1, 56) = 4.51, *p* = .038, *η_p_^2^*=.075) and between the two *Volatility* sessions (*F*(1, 56) = 11.18, *p* < .001, *η_p_^2^*=.167) in the first half of trials.

The analysis of Kalman gain *K_1_* revealed a significant main effect of *Volatility* in the first half of trials (*F*(1, 56) = 21.89, *p* < .001, *η_p_^2^*=.281): on average, participants placed 71.7% of the weight on the sensory input in the high-volatility session, but only 58.2% in the low-volatility session. Numerically, individuals with ASD placed greater weight on the sensory input (68%) than their TD counterparts (62%), but the difference did not reach significance (*Group*, *F*(1, 56) = 1.91, *p* = .172, *η_p_^2^* =.03. The interaction between *Group* and *Volatility* was also not significant (*F*(1, 56) = 0.02, *p* = .89, *η_p_^2^*<.001).

We then compared the first and second half of trials, which revealed significant changes of *K_1_* with *Volatility* (*F*(1, 56) = 6.18, *p* = .016, *η_p_^2^* = .099). Specifically, there was an increase of 2.6% in terms of the weight placed on the prediction error between the first and second half of trials in the high-volatility session and a decrease of 4.2% in the low-volatility session – a net difference of 6.8%. The change was comparable between the two groups, *F*(1, 56) = 0.75, *p* = .39, *η_p_^2^* = .013, indicating that both groups adjusted their weighting similarly.

Thus, the process of short-term belief updating remained relatively stable over the course of all trials, with the ASD group exhibiting a significantly higher Kalman gain *K_2_* relative to the TD group. This difference in *K_2_* influenced the rate at which long-term beliefs are formed, which in turn affected the central tendency effect. A significant difference in the central tendency effect between the two groups was evident in the first half of the trials, though it diminished across the entire session.

Last, we examined for potential relationships between the estimated parameters and symptom severity, measured by the Autism-Spectrum Quotient (AQ), Empathy Quotient (EQ), Systemizing Quotient (SQ), Beck’s Depression Inventory (BDI), and Empathising-systemising (E-S) scores (see Appendix 2). We only found a negative correlation between SQ and *K_2_* for both groups: individuals with higher SQ (that is, systematic-thinking) scores tended to exhibit lower updating weights *K_2_*. However, *K_2_* was significantly higher for individuals with ASD than their TD counterparts.

## Discussion

The nature of differences in predictive processing between individuals with ASD and their TD peers remains a subject of intense debate (Cannon et al., 2021; Lawson et al., 2014, 2017; Noel et al., 2022; Noel & Angelaki, 2023; Palmer et al., 2017; Pellicano & Burr, 2012; Sinha et al., 2014; Van de Cruys et al., 2014). Opinions are divided: some studies suggest that predictive processes are generally impaired in ASD (Karaminis et al., 2016; Lawson et al., 2014; Pellicano & Burr, 2012; Van de Cruys et al., 2014), others find that predictive processing remains intact (Angeletos Chrysaitis & Seriès, 2023; Noel et al., 2022; Pesthy et al., 2023; Randeniya et al., 2021; Van de Cruys et al., 2018), even though showing atypical learning dynamics (Lieder et al., 2019; Randeniya et al., 2021; Vishne et al., 2021). We hypothesized that this divergence in findings reflects differences in how the studies handled the updating of prior beliefs, perceptual estimates derived from prior knowledge and sensory input, and the dynamics across the timescales that they examined. Consistent with this, applying (Glasauer & Shi, 2022)) the two-state Bayesian model to performance in a duration-reproduction task under conditions of high vs. low cross-trial duration volatility, we distinguished differential updates of prior means and perceptual estimates between the ASD and the TD group, and how these updates influence long-term prediction and behavioral outputs.

In line with the application of Glasauer and Shi’s (2022) two-state Bayesian model to performance in a duration-reproduction task under conditions of high versus low cross-trial duration volatility, we distinguished the autistic spectrum disorder (ASD) group from the typically developing (TD) group. The differentiation was based on how each group updated the prior means and perceptual estimates, and how these updates influenced long-term prediction and behavioral outcomes.

Our results revealed both groups to exhibit similar central-tendency effects, in line with recent findings (Poole et al., 2022), and both to be responsive to changes in environmental uncertainty, evidenced by a more pronounced central tendency under high-relative to low-volatility conditions. The ASD group did *not* show an overestimation of the volatility, contrasting with previous findings (Lawson et al., 2017). Thus, according to our results, the primary challenge for individuals with ASD does not appear to lie in the formation of priors or in sensory prediction. In fact, participants in our ASD group proved entirely capable of forming appropriate priors, consistent with recent evidence from successful one-shot learning (Van de Cruys et al., 2018), learning from preceding trials (Pell et al., 2016), and adaptation to statistical settings (Allenmark et al., 2021; Hense et al., 2019; Pantelis & Kennedy, 2017).

Critically, however, compared to TD participants, we found individuals with ASD to rely more heavily on the sensory input specifically for the iterative updating of priors, pointing to a higher level of uncertainty in their prior mean distributions. In contrast to previous suggestions (e.g., Lawson et al., 2017; Van de Cruys et al., 2014), though, their increased reliance on sensory input for revising interim priors was not mirrored in a heightened reliance on sensory input for updating perceptual estimates: participants with ASD actually showed a similar weighting of sensory inputs (*K_1_*) in estimating durations compared to their TD counterparts. Yet importantly – and potentially a source of difference to the findings in previous studies –, our observations suggest that differences in *average* outcomes across different timescales may arise due to this distinct approach to updating priors: in the first half of trials within each session, the ASD group did exhibit a smaller central-tendency effect than the TD group. While being consistent with previous reports (Karaminis et al., 2016; Pellicano & Burr, 2012), this group difference diminished over the course of the session – showing that individuals with ASD did eventually acquire the sample distribution of durations like e TD participants.

The two-state Bayesian model (Glasauer & Shi, 2022) differentiates two types of updates: updating of the current perceptual estimate (Eq. 4) and iterative updating of the prior based on the recent past (Eq. 5). The fact that individuals with ASD placed more weight on sensory input when updating their interim priors implies that they perceived temporal events as less interrelated than their TD counterparts, leading them to adjust their beliefs more in response to the immediate sensory data. This process influences both their initial prior beliefs and the speed at which their priors are adapted to changing environmental conditions. Consequently, during the first half of each session, they showed a weaker central-tendency bias; yet by the end of the session, their performance was indistinguishable from that of their TD peers, in both the high- and low-volatility settings. This bolsters the idea that individuals with ASD can accurately gauge environmental statistics, but their inter-trial updating differs from that of TD controls (Allenmark et al., 2021).

To some extent, the specific atypicality in inter-trial prior updating that we observed aligns with recent suggestions that the primary atypicality in ASD involves a slower adaptation speed (Lieder et al., 2019; Vishne et al., 2021). For instance, Lieder et al. (2019), in a tone-discrimination task, found individuals with ASD to place smaller weights on the immediately preceding trials compared to controls, although their long-term weighting turned out similar. However, the atypical updating to which our study points is different: it is not the direct *integration* of past information into perceptual estimations that varied between groups, but rather the *updating* of the internal prior beliefs. To further rule out that the incorporation of recent history into duration estimates differed, we conducted a general linear model (GLM) (detailed in Appendix 3) that included both the current and the immediately preceding trial durations to predict the current reproduction errors. While this analysis revealed a significant difference in the weighting of the preceding duration between the high- and low-volatility sequences, there was no difference between the ASD and TD groups – suggesting that both groups similarly integrate preceding stimuli into duration judgments.

It is important to note that the distinctive updating of prior beliefs in ASD is not necessarily, or per se, a ‘slower’ process. Rather, it is characterized by a heavier reliance on sensory inputs, which might slow down the adaptation speed. In uncertain environments, this reliance on sensory input, rather than on prior beliefs, would actually be beneficial: it ensures a more responsive adaptation to the current stimuli, leading to a less pronounced central-tendency bias early on in the session. However, this approach yields a somewhat slower adaptation in the longer run (here, across the entire session). We speculate that this might be reflected in slower adaptation of behavior in everyday life, that is, behavioral rigidity.

The distinctive updating of prior beliefs that we observed in individuals with ASD may clarify some mixed results in the literature regarding predictive processing in ASD (Angeletos Chrysaitis & Seriès, 2023) – which often depend on the rate of adaptation and timescales used in the respective experiments. Thus, for example, some studies indicate that individuals with ASD maintain an intact sensitivity to interval timing (Isaksson et al., 2018; Jones et al., 2009; Mostofsky et al., 2000; Poole et al., 2022), whereas others report their sensitivity to be reduced, especially with certain intervals or tasks (Falter et al., 2012; Isaksson et al., 2018; Martin et al., 2010; Szelag et al., 2004). Notably, Karaminis et al. (2016) found children with ASD to exhibit a stronger central tendency and a poorer temporal resolution in duration reproduction than their TD counterparts. However, their central tendency was weaker than would be predicted by Bayesian integration, implying poorer priors compared to the control group. Interestingly, though, Karaminis et al. (2016) conducted their study with only 77 trials per whole session for child participants, fewer than the first half of trials within a session (i.e., 125 trials) in our study. Accordingly, their findings might reflect an initial low weight of prior beliefs in sensory-prior integration, consistent with our observations from the first halves of the sessions. One hypothesis deriving from our findings would be that extended testing might allow children with ASD to develop similar priors to their TD counterparts.

It is worth noting that within the ASD group, three individuals (out of our sample of 32) exhibited markedly different behavior from the rest, while also showing strikingly similar response patterns among themselves (see Appendix 1). In the relatively stable and predictable (i.e., the low-volatility) sequence, these individuals reproduced time durations proportional to the target durations (i.e., the actual stimulus), with some general over- or underestimations. However, in the high-volatility environment, they consistently reproduced the same duration across all sampled intervals, effectively ignoring the external sensory inputs (with *K_1_* and *K_2_* values at 0.07 and 0.04, respectively) and relying solely on an overly strong internal prior duration. The normal reproduction performance observed in the low-volatility sequence rules out that their ‘outlier’ behavior was simply due to a misunderstanding of the instructions. Interestingly, during the debriefing, these three individuals reported no awareness of the difference in the volatility regimen between the two sessions, suggesting that their performance was influenced implicitly. We hypothesize that when experiencing a highly volatile stimulus sequence, these participants responded by treating the large uncorrelated stimulus changes as noise instead of signal, thus effectively ‘discounting’ the sensory input. That is, they appear to change their model between high- and low-volatility environments concerning the likelihood distribution, the assumption about what is noise and what is signal. It should be noted that even when including these outliers in our analysis, the critical results of the two-state model remained consistent.

In conclusion, our findings argue in favor of a new perspective of predictive-processing dynamics in ASD, while helping to clarify apparent inconsistencies in the literature. We found that individuals with ASD use prior knowledge similarly to controls. But, critically, they place greater emphasis on sensory information specifically during the updating of interim priors (distinct from the updating of percepts). This distinctive process may lead to slower adaptation of perceptual decision-making across time, which, depending on the environmental volatility, may not necessarily be a disadvantage. Discrepant results in the extant literature might be due to the various studies focusing on different timescales and adaptation stages, and ignoring the dynamics of the two separable underlying updating processes.

Our observations indicate that individuals with ASD adapt appropriately to various environments, whether highly volatile or stable. However, as revealed by our two-state iterative updating model, they display a unique inter-trial dynamics pointing to a distinctive process of iterative prior updating. Accordingly, it is a focus on temporal discontinuity and overreliance on sensory input during moment-to-moment prior updating that are characteristic of predictive processing in ASD.

## Materials and Methods

### (a) Participants

32 individuals (13 females, 19 males, aged between 18 and 67 years, *M* = 32.0; *SD* = 12.3) with confirmed ICD-10 ASD diagnosis (World Health Organization, 1992) of F84.0 or F84.5 were recruited from the database and network partners of the Outpatient Clinic for Autism Spectrum Disorders at the Department of Psychiatry, LMU Munich, Munich, Germany. 32 TD controls (13 females, 19 males, aged between 18 and 70 years, *M* = 31.6*, SD =* 13.6) with no reported history of mental illnesses or neurological deficits were recruited via local advertising. The groups were matched pairwise using the ‘Wortschatztest’ (WST) (Schmidt & Metzler, 1992), a measure of crystalline intelligence. Both groups completed the Autism-Spectrum Quotient (AQ) (Baron-Cohen et al., 2001), Empathy Quotient (EQ) (Baron-Cohen & Wheelwright, 2004), Systemizing Quotient (SQ) (Baron-Cohen et al., 2003), and Beck’s Depression Inventory (BDI) (Beck et al., 2011) as proxies of symptom load. The groups were comparable in terms of IQ (*p*=.28), and age (*p*=.9), but differed significantly on the AQ (*p* < .001), EQ (*p* < .001), SQ (*p* = .005), and BDI (*p*=.018) scores. In addition, the empathizing-systemizing (E-S) score (given by the difference between the EQ and SQ) also showed a significant difference (*p* < .001) (see Appendix 3, Table A2).

All participants provided written informed consent prior to the experiment and received a compensation of 10 Euros per hour for their service. The study was approved by the Ethics Board of the LMU Munich Faculty of Pedagogics and Psychology.

### (b) Design and procedure

The experiment was conducted in a sound-attenuated and moderately lit experimental cabin. A visual stimulus in the form of a filled-in yellow disk patch (diameter: 4.7° of visual angle; luminance: 21.7 cd/m^2^) was used for delivering (stimulus) durations on a 21-inch LACIE CRT monitor (refresh rate: 85 Hz). The experimental code was developed using the Matlab PsychToolbox (Kleiner et al., 2007).

We adopted the duration-reproduction paradigm (Ren et al., 2021) (Figure 1a). A trial started with a central fixation cross (size: 0.75° degrees of visual angle) presented for 500 ms. This was followed by a white dot (diameter: 0.2°) in the center, signaling participants to press and hold either the left or the right mouse button to initiate the duration-encoding phase. Once the button was pressed, a yellow disk appeared on the screen for a randomly chosen duration ranging from 100 ms to 1900 ms (elaborated in the next paragraph). The yellow disk then disappeared, prompting the participant to immediately release the mouse button. A blank screen lasting 500 ms separated the duration-encoding and -reproduction phases. Next, a white dot appeared, telling participants to reproduce the duration they had just observed by pressing and holding the mouse button for a duration matching that of the previously displayed yellow disk, and then releasing the button. Upon pressing the mouse button, the yellow disk reappeared in the display center, and it disappeared once the button was released. Subsequently, a feedback display was presented for 500 ms, informing participants about the accuracy of their reproduction. The feedback display consisted of five horizontally arranged disks graphically indicating the ratio of the reproduction error relative to the actual duration. From left to right, the disks denoted the following relative error ranges: less than −30%; between −30% and −5%, between −5% and 5%, between 5% and 30%, and greater than 30%. One of the disks was highlighted (i.e., changed its color from the initial white) – either in green for one of the three central circles or in red for one of the outer circles –, indicating the error range on a given trial. The red highlighting signaled a large duration under- or, respectively, overestimation, that was to be avoided.

The experiment was divided into two sessions, each comprising 10 blocks of 25 trials (250 trials in total), with one session presenting a low- and the other a high-volatility sequence of stimulus durations. Critically, both conditions consisted of the same set of stimulus durations and an equal number of duration repetitions; they differed only in terms of the presentation order. First, individually for each participant (i.e., matched pair of participants), a low-volatility sequence of durations was generated using a random-walk process that began with a randomly selected duration. The duration of each subsequent trial was determined based on the duration of the immediately preceding trial to which a small random fluctuation, sampled from a normal distribution, was added. Over the course of 250 trials, these fluctuations could (occasionally) accumulate and exceed the pre-set range (e.g., yielding a negative value). To avoid this, the durations were scaled back to fit within 100 ms to 1900 ms and then rounded to the nearest 100-ms value, allowing for multiple repetitions of the tested durations. This procedure resulted in a ‘low-volatility’ sequence characterized by mild fluctuations across trials. To create the ‘high-volatility sequence’, we randomly shuffled the durations from the low-volatility sequence, producing greater trial-to-trial variations. Figure 1b illustrates typical sequences for the low- and high-volatility sequences across two consecutive sessions. Both sequences were generated prior to the experiment, and the order of the low- and high-volatility sessions was counterbalanced across participants.

Importantly, we administered identical sequences to (age- and IQ-) matched participants in the ASD and TD groups, ensuring that any differential effects observed would be attributable to differences between the two groups.

Prior to the experiment, participants received comprehensive written and verbal instructions, and they underwent a pre-experimental training block of at least 10 trials to ensure they understood the instructions. Subsequently, the experiment proper was conducted, followed by filling-out of the questionnaires.

#### Ethic

The study was approved by the Ethics Board of the Faculty of Pedagogics and Psychology at LMU Munich, Germany.

## Data accessibility

Experimental codes, data, and analyses are published at g-node: 10.12751/g-node.85py1h.

## Acknowledgments

The study was supported by the German-Science-Foundation (DFG) research grant SH 166/3-2 awarded to ZS. CFW was supported by an excellence stipend from the Bavarian Gender Equality Grant (BGF) and DFG grants FA 876/3-1 and FA 876/5-1. SG was supported by DFG grant GL 342/3-2.

## Appendix

### Appendix 1. Duration reproduction in the high-(vs. low-) volatility session by three ‘outlier’ individuals in the ASD group

In the high-volatility session, three participants from the ASD group exhibited a markedly different performance compared to the others in the group (see Figure 3 Boxplots). To better visualize their reproduction performance, we replot their reproduced durations as a function of the physical duration. Figure S1 illustrates their performance in the low- and high-volatility sessions. As can be seen, the reproduced durations were similar across the range of durations in the high-volatility session (orange, ‘flat’), even though these individuals performed similarly to their peers in the low-volatility session. For these three outliers, the central-tendency indices (1 – regression coefficient) were extremely high (0.904, 0.997, and 0.907, respectively) in the high-volatility session. This suggests that these individuals relied heavily on their prior instead of the sensory inputs for their duration judgments, rendering a remarkable level of rigidity in their responses.

**Figure S1.**
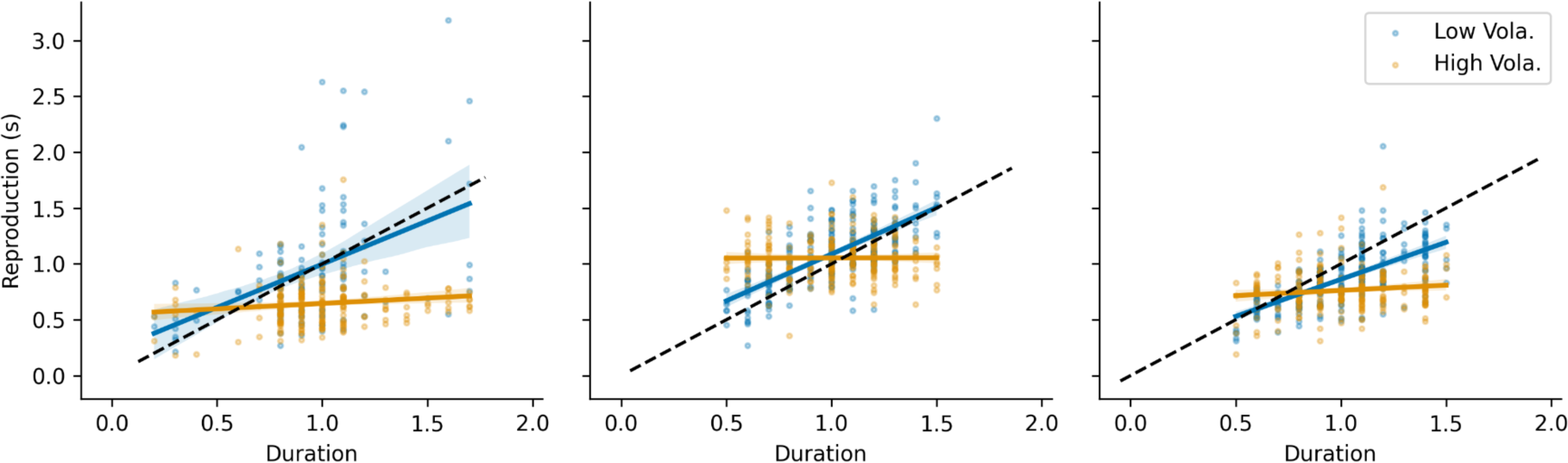
Duration reproduction by three ‘extreme’ ASD individuals, separately for the high-(orange) and low-volatility (blue) sessions. The diagonal dashed line denotes veridical reproduction.

### Appendix 2. Correlations between the updating weight and symptom severity

We examined for potential relationships between the estimated *K_2_* parameter and symptom severity, measured by the Autism-Spectrum Quotient (AQ), Empathy Quotient (EQ), Systemizing Quotient (SQ), Beck’s Depression Inventory (BDI), and Empathising-systemising (E-S) scores. In addition, we considered potential contributions from the two experimental factors *Group* and *Volatility*. We found a strong negative correlation between the *K_2_* and SQ scores (*p* = .003), and a strong positive correlation between the *K_2_* and E-S (*p* = .016) scores. The estimated linear models were:

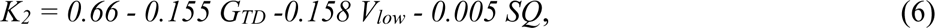

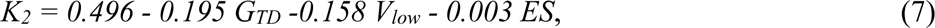

where *G_TD_* is a dummy variable for Group (1 for the TD group, 0 for the ASD group) and *V_low_* for the Volatility (1 for the low-volatility session, and 0 for the high-volatility session). The intercepts and coefficients for the dummy variables were all significant (*p* < .001). *K_2_* was significantly higher for the ASD group compared to the TD group (15.5% and 19.5%, respectively, in the above two estimated models). Figure S2 shows the correlation trends for both scores. Given that the E-S scores were derived from the difference between the Systemizing Quotient (SQ) and Empathy Quotient (EQ) scores, SQ is likely the main driving factor of the correlation.

The correlation suggests that the updating weight *K_2_* is sensitive to the individual preference and aptitude for systemizing: the higher the systemizing preference, the smaller the updating weight of the sensory data in the formation of the prior beliefs.

**Figure S2.**
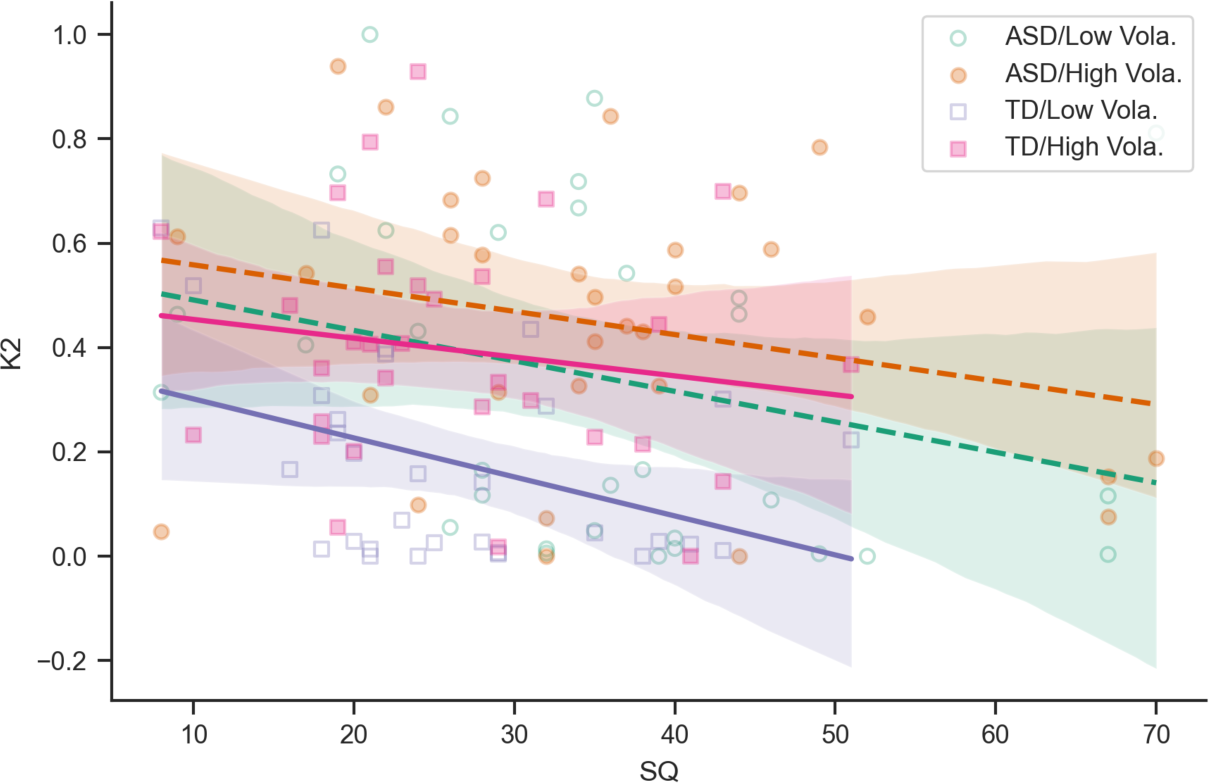
Negative correlations between the *K_2_* and Systemizing-Quotient (SQ) scores, separately for the ASD/Low-volatility (green circles, dashed line) and ASD/High-volatility (red circles, dashed line), TD/Low-volatility (light blue squares, solid blue line), TD/High-volatility conditions (pink squares, pink solid line).

### Appendix 3. A general linear model (GLM) considering the preceding trial

To examine whether there were any differences in how the two groups incorporated short-term trial history, we conducted a general linear model (GLM) that factored in both the current and the immediately preceding durations to predict the current reproduction errors. Given that in the low-volatility session, the current and previous durations were not fully independent, we used the duration difference of the current from the preceding trial (*ΔD = D_n_ – D_n-1_*) in our regression model. Additionally, to mitigate the impact of extreme performance, we removed the three ‘outlier’ individuals and their matched counterparts from the analysis. Table A1 displays the average regression coefficients and their associated standard errors.

We then performed separate mixed-design ANOVAs for each regression coefficient, with *Group* and *Volatility* as factors. These analyses failed to reveal any significant *Group* or *Volatility* effects on the intercept. The coefficients for the Current Duration *D_n_* differed significantly between the high- and low-volatility sessions, *F*(1,56) = 24.57, *p* < .001, *η_p_*^2^=.305. However, the main effect of Group and the Group x Volatility interaction failed to reach significance, *F*s < 2.218, *p*s > .14, *η_p_*^2^s< .038. Similarly, the coefficients for the deviation *ΔD* differed significantly between the high- and low-volatility sessions, *F*(1,56) = 54.49, *p* < .001, *η_p_*^2^=.489; but again, there were no (main or interaction) effects involving *Group*, *F*s < 3.34, *p*s > .073, *η_p_*^2^s < .056. We take this to indicate that the ‘history’ impact of the preceding Duration was similar for both groups.

**Table A1.** Average regression coefficients and their associated standard errors (in parentheses) for the ASD and TD groups, separately for the high- and low-volatility sessions.

| Group | Volatility | Intercept | Current Duration ($D_n$) | $D_n - D_{n-1}$ |
| --- | --- | --- | --- | --- |
| ASD | High | 0.032 (0.015) | -0.166 (0.026) | -0.104 (0.016) |
|  | Low | 0.042 (0.015) | -0.092 (0.030) | -0.281 (0.041) |
| TD | High | 0.029 (0.011) | -0.224 (0.023) | -0.123 (0.012) |
|  | Low | 0.038 (0.013) | -0.086 (0.021) | -0.394 (0.049) |

### Appendix 4: Descriptive characteristics of the ASD and TD groups

The groups were matched pairwise using the ‘Wortschatztest’(WST, Schmidt & Metzler, 1992), a measure of crystalline intelligence. Both groups completed the Autism-Spectrum Quotient (AQ, Baron-Cohen et al., 2001), Empathy Quotient (EQ, Baron-Cohen & Wheelwright, 2004), Systemizing Quotient (SQ, Baron-Cohen et al., 2003), and Beck’s Depression Inventory (BDI, Beck et al., 2011). The mean scores and paired *t*-tests are shown in Table A2. As can be seen, the AQ, EQ, SQ, and BDI scores were significantly different between the two groups.

**Table A2.** Descriptive characteristics (Means and SDs) for ASD and TD group.

| *Measures* | *ASD (n=32)* | *TD (n=32)* | *Group comparison* |
| --- | --- | --- | --- |
| Age | 32.03 (12.3) | 31.6 (13.6) | $t_{31} = 0.13, p = .898$ |
| IQ score | 107.4 (9.7) | 109.1 (13.4) | $t_{31} = 0.573, p = .28$ |
| Autism-Spectrum Quotient score | 36.7 (7.4) | 15.8 (6.0) | $t_{31} = 18.06, p < .001^{**}$ |
| Empathy Quotient score | 26.97 (12.32) | 50.38 (14.46) | $t_{31} = -7.85, p < .001^{**}$ |
| Systemizing Quotient score | 35.25 (14.93) | 26.63 (10.1) | $t_{31} = 3.05, p < .01^{**}$ |
| Beck's Depression Inventory score | 10.69 (8.37) | 5.25 (7.46) | $t_{31} = 2.49, p = .018^*$ |

Note: \*\* denotes $p < .05$ and \*\*\* $p < .001$ .

## Footnotes

1 The likelihood distribution is assigned to the single measurement to represent the uncertainty of the measurement. Accordingly, the likelihood distribution represents the measurement together with its uncertainty (which is assumed to be known to the observer).

## Notes

https://gin.g-node.org/msense/predictive_coding_in_asd

### Competing Interest Statement

The authors have declared no competing interest.

### Summary of Updates

This revision refined the two-state model used in the study and update the illustration of the model for better understanding.

